# Siderophore-Mediated Conveyance of Antibacterial-Antisense Oligomers

**DOI:** 10.1101/2023.10.30.564798

**Authors:** Mathijs J. Pals, Luuk Wijnberg, Çağlar Yildiz, Willem A. Velema

## Abstract

Antibacterial resistance is a major threat for human health. There is a need for new antibacterials to stay ahead of constantly-evolving resistant bacteria. Antibiotic antisense oligomers hold promise as powerful next-generation antibiotics, but issues with their delivery hamper their applicability. Here, we exploit the siderophore-mediated iron uptake pathway to efficiently transport antisense oligomers into bacteria. We appended a synthetic siderophore to antisense oligomers targeting the essential *acpP* gene in Escherichia coli. Siderophore-conjugated morpholino and PNA antisense oligomers displayed potent antibacterial properties. Conjugates bearing a minimal siderophore consisting of a mono-catechol group showed equally effective. Targeting the *lacZ* transcript resulted in dose-dependent decreased β-galactosidase production, demonstrating selective protein downregulation. Whole-genome sequencing of resistant mutants and competition experiments with the endogenous siderophore verified selective uptake through the siderophore-mediated iron uptake pathway. Lastly, no toxicity towards mammalian cells was found. Collectively, our work provides a convenient approach for delivering antisense oligomers into bacteria.

## Introduction

The alarming rise in antibiotic-resistant bacteria poses a major threat for human health.^1,2^ Increasing reports of pan-drug resistant pathogens are a forewarning of a future in which infections are untreatable. It is estimated that by 2050, 10 million annual deaths will be caused by drug-resistant bacteria.^3^ To counteract this frightening prospect we need to restock our arsenal of effective antimicrobials.^3,4^ Despite increased awareness, few new antibiotics have reached the market in the last decades and there is a clear need for advanced therapies and approaches.^5^

One promising development in the search for new antibiotics is antibacterial antisense oligomers (ASOs).^6–8^ ASOs are short, often chemically modified, nucleic acid sequences that can hybridize to a complementary mRNA target (**Fig. 1a**).^6,9,10^ Hybridization blocks the ribosome from binding to the mRNA and can cause mRNA degradation by nucleases depending on the nature of the ASO (**Fig. 1a**).^6,9–11^ Since the ASO sequence can be programmed, any mRNA of interest can be targeted rendering this approach extraordinarily powerful and highly promising for next-generation therapies including antibiotic development. Indeed, numerous studies have shown that ASOs targeting essential genes are effective in inhibiting bacterial growth.^12–16^ However, the major limiting factor of ASOs as antibacterial agents is transport of these large molecules into the bacterial cell, so far hindering their clinical application.^17,18^

**Figure 1:**
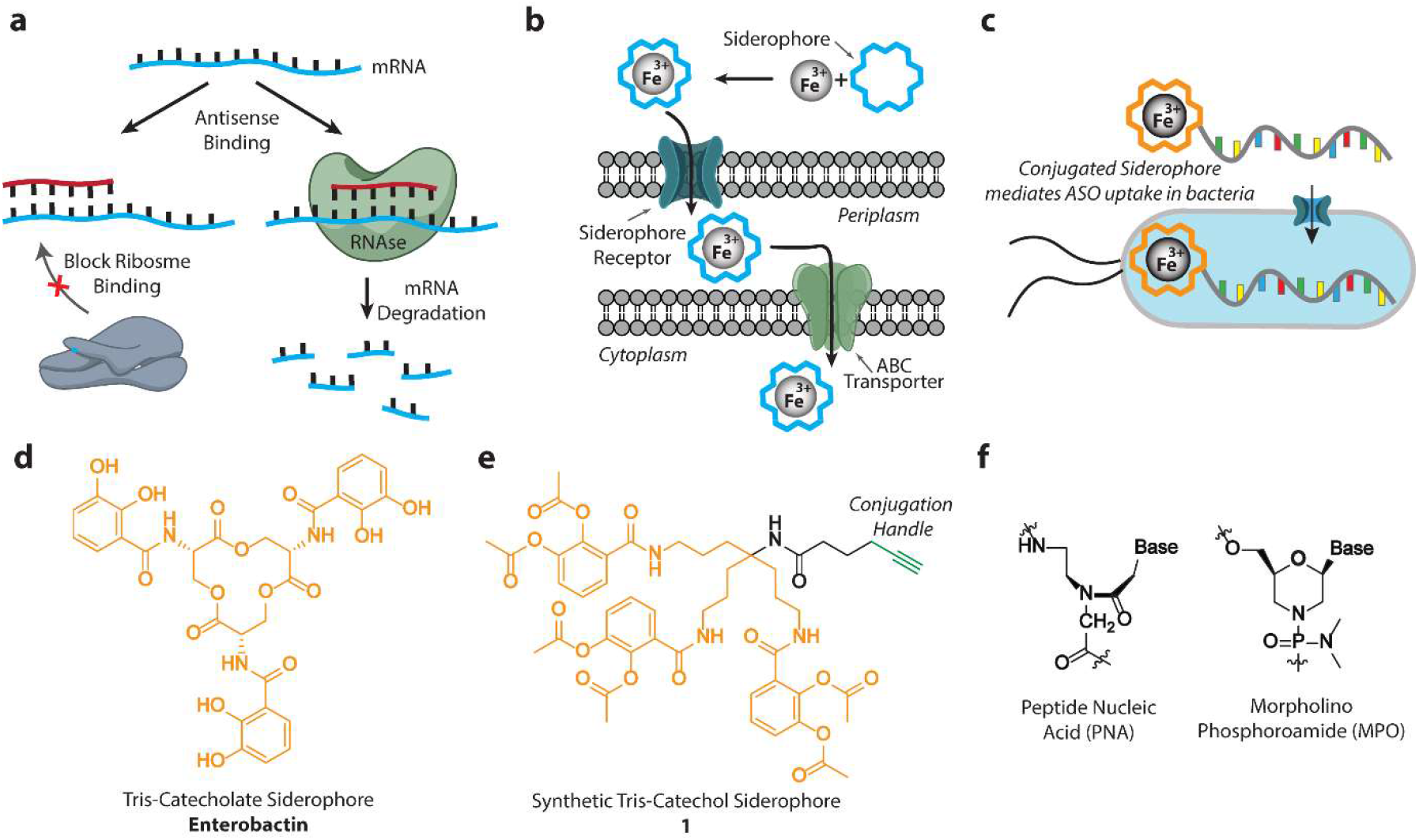
Design mechanism and uptake of antibacterial ASO-siderophore conjugates (a) Antisense mechanism. (b) Siderophore uptake route. (c) Siderophore-mediated ASO uptake in bacteria. (d) Molecular structure of enterobactin. (e) Molecular structure of ‘clickable’ siderophore **1**. (f) Molecular structures of PNA and MPO-based ASOs.

The most frequently used delivery system for antibacterial ASOs is cell-penetrating peptides (CPPs). These short cationic or amphiphilic peptides can be appended to the ASO to increase bacterial uptake via a not fully understood penetration mechanism.^19^ Although capable of transporting ASOs into bacteria, CPPs can be cytotoxic by themselves resulting in severe toxicity in mammals.^20–22^ Moreover, CPPs do not selectively penetrate bacterial cells, but mammalian cells as well.^23^ This results in potential additional off-target effects of the ASO on human mRNAs limiting their clinical use.^24^ Given these disadvantages of CPPs, there is a need for selective, non-toxic and versatile carrier molecules for ASO-delivery.

Selective delivery in bacteria could be secured by hijacking a transport system that is not found in humans, such as the siderophore-mediated iron uptake pathway.^25^ Siderophores are small Fe(III) chelating molecules synthesized by bacteria.^26^ After excretion of the siderophore it chelates Fe(III) and the subsequent complexes are actively taken up by bacterial cells via receptor mediated transport (**Fig. 1b**).^25^ It has been shown that this transport route can be hijacked to convey small-molecule cargo into bacteria, including small-molecule antibiotics, by appending a (synthetic) siderophore to the cargo.^27–33^ We hypothesized that this Trojan Horse approach might be applicable to larger cargo as well, such as antibacterial ASOs and could be exploited to devise potent and selective antibacterial antisense agents (**Fig. 1c**).

In this article, we report the potential power of combining the selectivity and programmability of antibacterial antisense technology with siderophore-mediated conveyance. Synthetic catechol siderophores (**Fig. 1d, e**) were conjugated to ASOs targeting the essential *acpP* gene in the ESKAPEE pathogen *Escherichia coli* (*E. coli*). The ASO-siderophore conjugates displayed potent micromolar antibacterial properties compared to the naked antisense oligomers which did not affect bacterial growth up to 100 μM. Siderophore-mediated conveyance was successful for the two commonly used antisense backbones: peptide nucleic acids (PNA), and morpholino phosphoroamide (MPO).^13–15^ Selective downregulation of target gene products was demonstrated by targeting the *lacZ* gene. In viability studies, no cytotoxicity of the ASO-siderophore conjugates on human cells was observed. Finally, to demonstrate that the uptake mechanism is through the siderophore-mediated iron transport pathway, competition experiments with the endogenous enterobactin siderophore were performed and resistant mutants were generated and analyzed using PacBio whole genome sequencing. Mutations were pinpointed to an SNP resulting in translation termination of the *ybiX* gene, which forms an operon with the outer membrane iron-catecholate transporter and is a suspected iron-uptake factor. ^34^

## Results and discussion

### Design and synthesis of siderophore-ASO conjugates

To explore the potential of siderophore mediated ASO delivery, we focused on the ESKAPEE pathogen *E. coli* since antisense delivery into this gram-negative bacterium is a major issue.^35,36^ The main siderophore of *E. coli* is enterobactin (**Fig. 1d**), which belongs to the catecholate class of siderophores.^25^ Recent studies with synthetic variants of enterobactin showed successful transportation of small cargo into the bacterial cell, making us hopeful that larger cargo such as ASOs could be transported as well.^37–39^ Inspired by pioneering work by Miller and coworkers,^40^ we synthesized an alkyne bearing synthetic tris-catechol siderophore (**Fig. 1e**) that can be appended to azide containing ASOs using a copper-catalyzed azide-alkyne cycloaddition (CuAAC).

Antisense oligonucleotides for antibacterial applications generally consist of chemically modified nucleic acids such as peptide nucleic acids (PNAs) and morpholino oligomers (MPO).^12,15,41,42^ Although similar in mode of action, PNAs and MPOs are structurally different (**Fig. 1f**) resulting in unique properties for each backbone. PNAs have been reported to be more flexible than MPOs but have poorer water-solubility.^43^ Nielsen and coworkers. showed that a minimum of 10-nt is required for sufficient selectivity and efficacy.^12^ Therefore, we chose a 10-nt PNA backbone as the starting point of the ASO (**Fig. 2a**). To improve the solubility of the PNA and generate space between the ASO and siderophore, we incorporated 2 hydrophilic PEG2 linkers at the N-terminus (or 5’) side. To enable conjugation with the siderophore by click chemistry, L-azidolysine was incorporated at the N-terminus (**Fig. 2b**).

**Figure 2:**
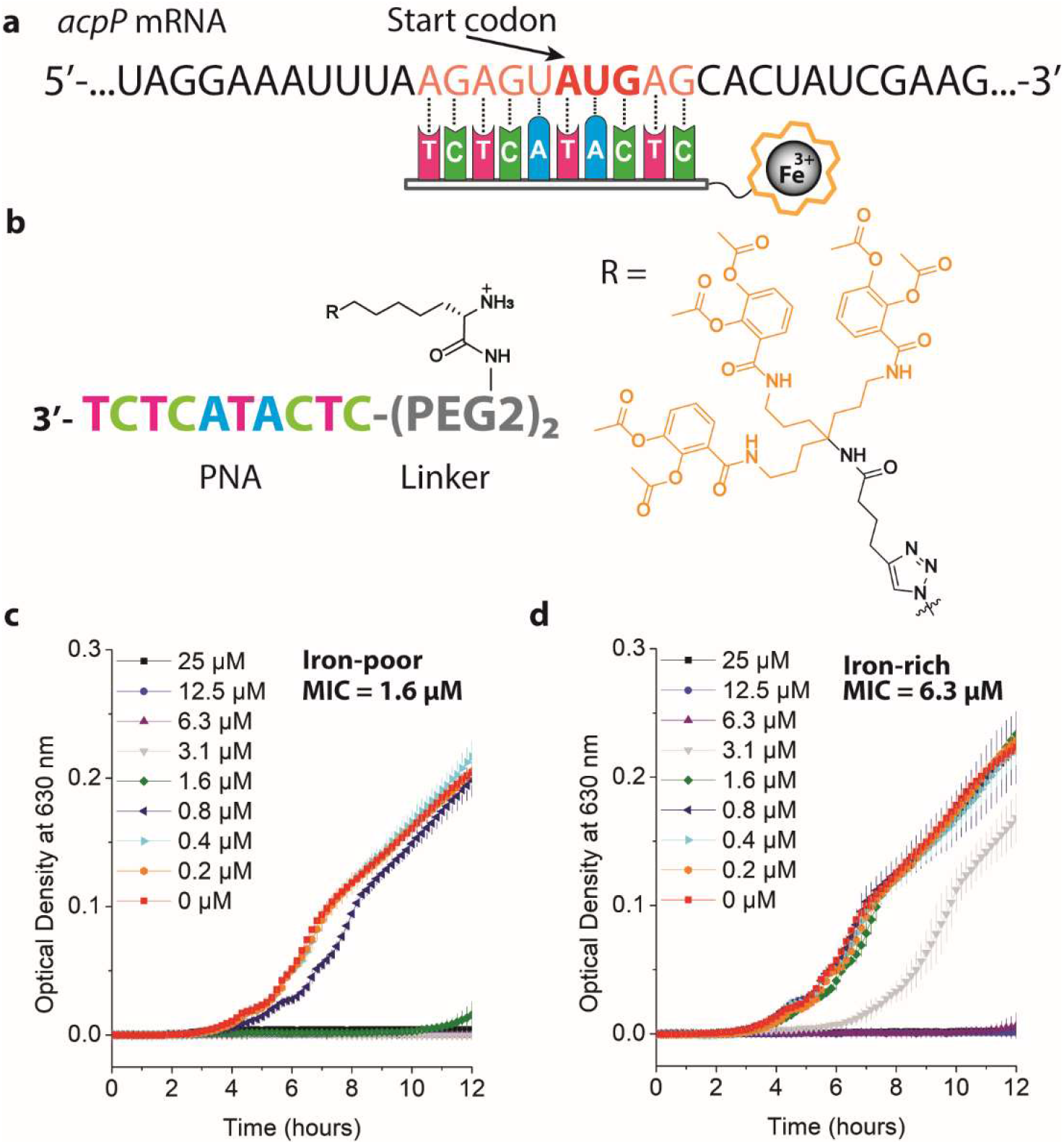
Bacterial-growth inhibition of antibacterial ASO-siderophore conjugate (a) Hybridization of the *acpP* PNA with the target mRNA. (b) Schematic representation of the siderophore-PNA conjugate (**ASO 2**) structure. (c) Growth of *E. coli* in presence of the siderophore-PNA conjugate (**ASO 2**) in iron-poor MH2 medium. (d) Growth *of E. coli* in presence of the siderophore-PNA conjugate (**ASO 2**) in iron-rich MH2 medium. Error-bars represent standard deviations based on three technical replicates.

### Inhibition of *E. coli* growth

To test the concept of siderophore-mediated delivery of antibacterial ASOs, we synthesized a known 10-nt lethal sequence that targets the well-studied essential *acpP* gene (**ASO 1**).^12,42,44,45^ This gene encodes an acyl carrier protein that is essential for fatty acid synthesis in *E. coli*.^12,44^ To test the antibacterial effect of the anti *acpP* PNA-siderophore conjugate (**ASO 2**), we performed standard minimal inhibitory concentration (MIC) experiments in Mueller Hinton 2 broth (MH2). The uptake of siderophores and sideromycins is increased in iron-poor conditions.^46^ Therefore, the MH2 medium was made iron-poor or iron-rich by addition of 2,2’-bipyridine or FeCl^3^ respectively. The bacteria were grown in iron-poor or rich MH2 at 37 °C with increasing concentrations of the anti *acpP* PNA-siderophore conjugate (**ASO 2**) for 12 hours. Gratifyingly, the conjugate inhibited growth of wild-type *E. coli* K12 with a MIC of 1.6 μM in iron-poor conditions (**Fig. 2c**). This observed growth inhibition is comparable to the CPP-PNA conjugate of the same sequence, which implies efficient uptake of the conjugate.^12^ Moreover, the MIC in iron-rich medium is 4-fold higher, indicating that the conjugate is likely taken up via the iron-transport route (**Fig. 2d**).

### Siderophore dependence and sequence selectivity

To demonstrate that growth inhibition is a result of the ASO-siderophore combination, we tested the antibacterial effect of the siderophore **1** only (without PNA) and the naked PNA (**ASO 1**) (without siderophore). The naked PNA (**ASO 1**) did not affect bacterial growth at any of the concentrations measured (**Fig. 3a, b**), resulting in a >60-fold higher MIC than the conjugate. Moreover, siderophore **1** itself did not display any inhibition of growth up to 100 μM as well (**Fig. 3c, d**). These results show that the ASO only inhibits bacterial growth when it is conjugated to the siderophore. To establish that the antibacterial effect is a likely result of translation inhibition of the target sequence, we synthesized a scrambled version of the anti *acpP* PNA (**ASO 3**) (**Fig. 3e**). This scrambled version has the same length and number of each nucleobase, but with limited complementarity. When conjugated to the synthetic siderophore, bacterial growth was observed at all concentrations tested (MIC >25 μM), with minor effect on growth displayed at high concentrations, potentially caused by interacting with other RNA targets. These results confirm that the observed inhibition of growth with the *acpP* targeting conjugate (**ASO 2**) is likely due to binding of the ASO to the mRNA target. ^45^

**Figure 3:**
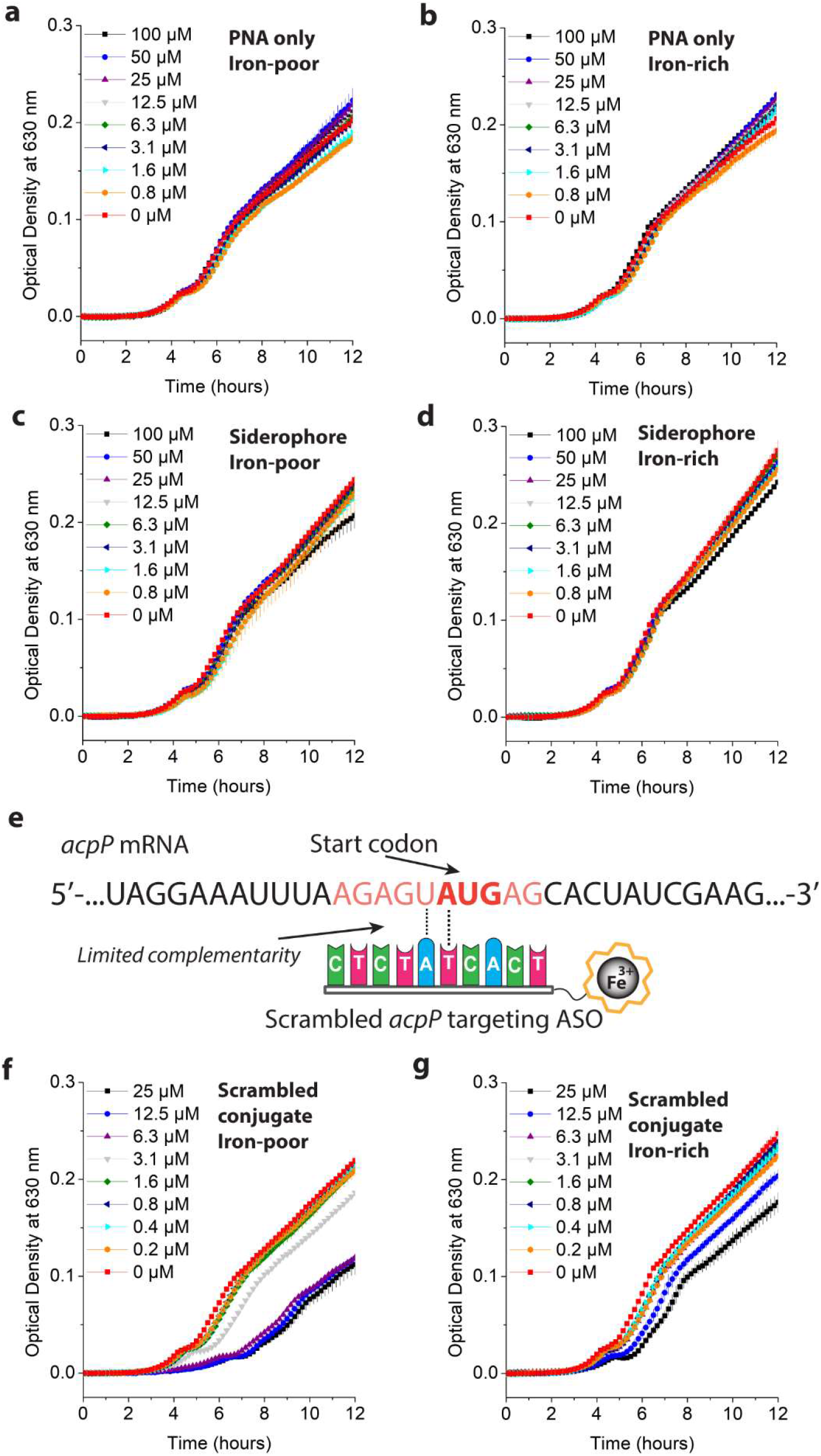
Effect of naked antibacterial ASO, siderophore **1** and scrambled ASO on bacterial growth. (a) Growth of *E. coli* in presence of the anti *acpP* PNA (**ASO 1**) only in iron-poor MH2 medium. (b) Growth of *E. coli* in presence of the anti *acpP* PNA (**ASO 1**) only in iron-rich MH2 medium. (c) Growth of *E. coli* in presence of the siderophore **1** only in iron-poor MH2 medium. (d) Growth of *E. coli* in presence of the siderophore **1** only in rich-poor MH2 medium. (d) Sequence of the non-complementary scrambled *acpP* PNA-siderophore conjugate (**ASO 4**). (f) Growth of *E. coli* in presence of the scrambled PNA conjugate (**ASO 4**) in iron-poor MH2 medium. (g) Growth of *E. coli* in presence of the scrambled PNA conjugate (**ASO 4**) in iron-rich MH2 medium. Error-bars represent standard deviations based on three technical replicates.

### Quantification of translation inhibition

To demonstrate that the antisense-based inhibition of translation results in down regulation of protein production, we synthesized an ASO-siderophore conjugate that targets a non-lethal gene with a conveniently quantifiable gene-product. We synthesized a 10-nt anti-*lacZ* PNA-siderophore conjugate (**ASO 6**) that should result in downregulation of its gene product β-galactosidase (**Fig. 4a**).^12^ To quantify the translation inhibition of the *lacZ* gene by this conjugate, *E. coli* were grown in iron-poor MH2 supplemented with 50 μM IPTG and increasing concentrations of the conjugate (**ASO 6**) for 4 hours at 37°C. Subsequently, the bacteria were permeabilized and the activity of β-galactosidase, was quantified (see **SI** for details).^12^ As expected, the matching anti *lacZ* PNA-siderophore conjugate (**ASO 6**) clearly inhibited the translation of its mRNA target, resulting in a decrease in relative β-galactosidase activity with an EC_50_-value of ∼1 μM (**Fig. 4b**). As a control, we synthesized an ASO-siderophore conjugate with a scrambled sequence (**ASO 8**), which did not result in a change in β-galactosidase activity (**Fig. 4c**). These results indicate that the ASO-siderophore conjugates likely display their observed antibacterial effect through inhibition of translation.

**Figure 4:**
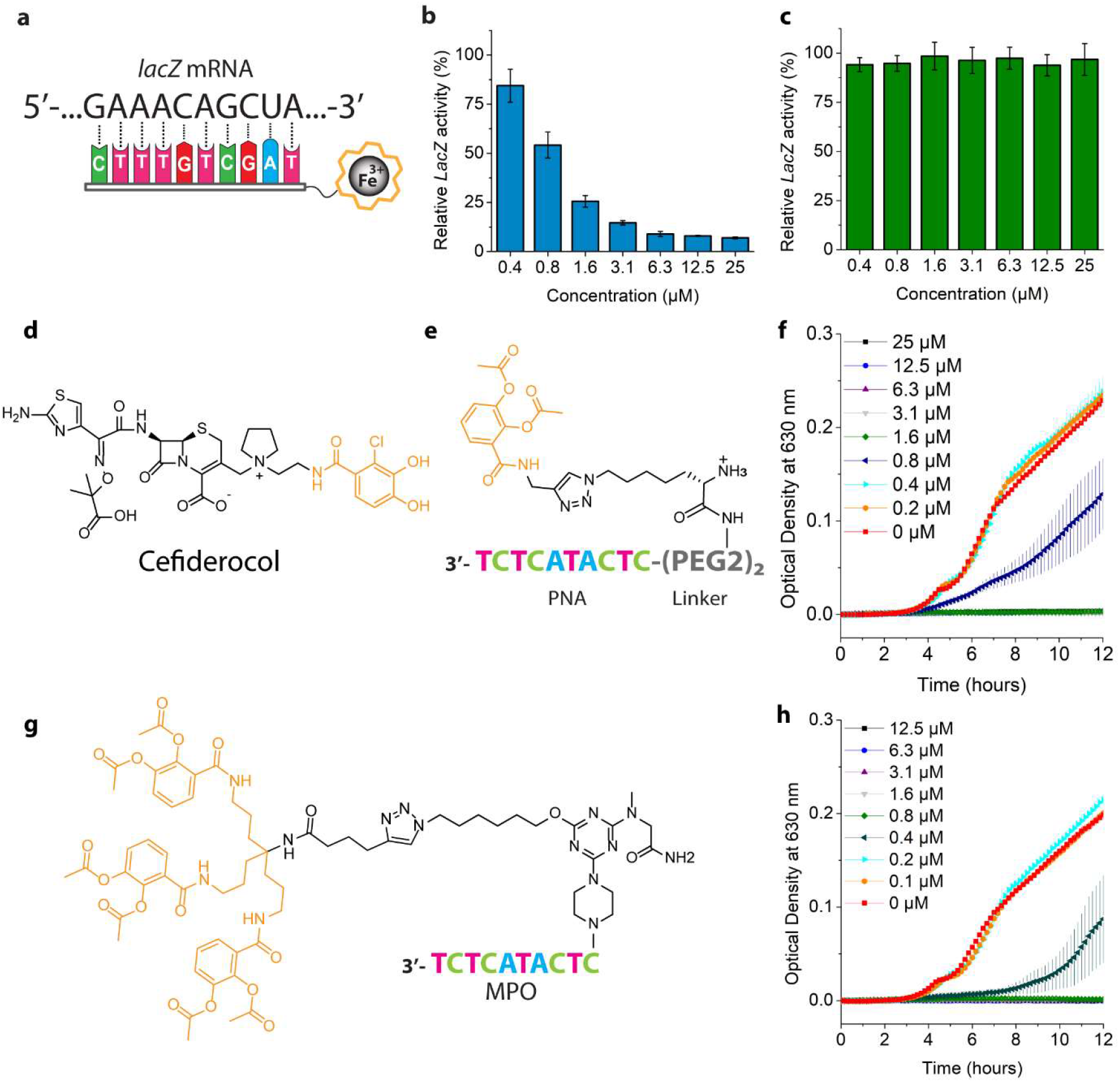
Effect on protein production and assessment of mono-catechol and MPO conjugates (a) Sequences of the targeted *lacZ* mRNA and PNA (b) Inhibition of *lacZ* translation by the anti *lacZ*-conjugate (**ASO 6**). (c) Inhibition of *lacZ* translation by the scrambled *lacZ* conjugate (**ASO 8**). (d) Chemical structure of Cefiderocol. (e) Schematic structure of the mono catechol-PNA conjugate (**ASO 9**) targeting the *acpP* gene. (f) Antibacterial effect of the mono catechol-PNA conjugate (**ASO 9**) on *E. coli* in iron-poor MH2. (g) Schematic representation of the siderophore-MPO conjugate (**ASO 11**). (h) Growth of *E. coli* in presence of the siderophore-MPO conjugate (**ASO 11**) in iron-poor MH2 medium. Error-bars represent standard deviations based on three technical replicates.

### Minimal structural requirements for siderophore-mediated uptake

The natural siderophore enterobactin, bears three catechol groups. However, several studies have shown that a single catechol group can be sufficient for siderophore-mediated uptake.^28,30,38^ This is illustrated by the clinically approved small-molecule antibiotic Cefiderocol, bearing a single catechol siderophore group (**Fig 4d**) that enhances antibiotic uptake.^47^ To test the minimal structural requirements for siderophore-mediated uptake of PNAs, we synthesized a single catechol group bearing an alkyne (siderophore **2**) and coupled this to the 10-nt *acpP* PNA (**ASO 1**) and tested its activity in MIC studies (**Fig. 4e**). Remarkably, the mono-catechol conjugate (**ASO 9**) displayed clear inhibition of growth with an MIC-value of 1.6 μM (**Fig. 4f**), while the mono-catechol siderophore **2** only did not show any antibacterial effect. These surprising findings indicate that appending a small catechol group to an ASO of >15x its molecular weight, results in efficient delivery. This highlights the simplicity and power of siderophore-mediated conveyance of ASOs.

### Morpholino-based conjugates

MPO- and PNA-based oligos are both used as antibacterial ASOs, but have significant structural differences (**Fig. 1f**). To test if siderophores can transport MPOs into bacteria as well, we conjugated the tris-catechol siderophore to an MPO with the same 10-nt anti *acpP* sequence as for the PNA version (**Fig. 4g**). Interestingly, the MPO-siderophore conjugate (**ASO 11**) showed a slightly increased antibacterial effect as compared to PNA with an MIC of 0.8 μM in iron-poor medium (**Fig. 4h**). As a control, the naked MPO sequence (**ASO 10**) was tested on antibacterial properties and no growth inhibition was observed. These results show that the siderophore-mediated transport of ASOs is not limited to PNAs only and can be applied to MPOs as well.

### ASO conveyance is through the siderophore-mediated iron uptake pathway

The observed result that the anti *acpP* PNA-siderophore conjugate shows a lower MIC in iron-poor than iron-rich medium indicates that the transport is likely due to the iron-uptake pathway. To further demonstrate this, we performed a competition experiment with the endogenous siderophore, enterobactin. It is expected that the endogenous siderophore will compete with the synthetic conjugate, lowering its uptake. To this end, the MIC assay of the anti *acpP* PNA-siderophore conjugate (**ASO 2**) was performed in presence of 10 μM of enterobactin and the optical density at 630 nm was measured over time. Interestingly, the presence of 10 μM enterobactin led to a strong decrease in the antibacterial effect resulting in a MIC >12.5 μM (**Fig. 5a**). These results are in line with similar competition experiments of sideromycins and endogenous siderophores and confirm that the conjugate is likely taken up via the enterobactin-route.^48,49^ We do not expect that competition with the endogenous siderophore will be problematic for the antibacterial application of the ASO-siderophore conjugates, since these were tested on enterobactin-producing wild-type *E. coli* K12 and showed good antibacterial effect with a low micromolar MIC-value (1.6 μM).

**Figure 5:**
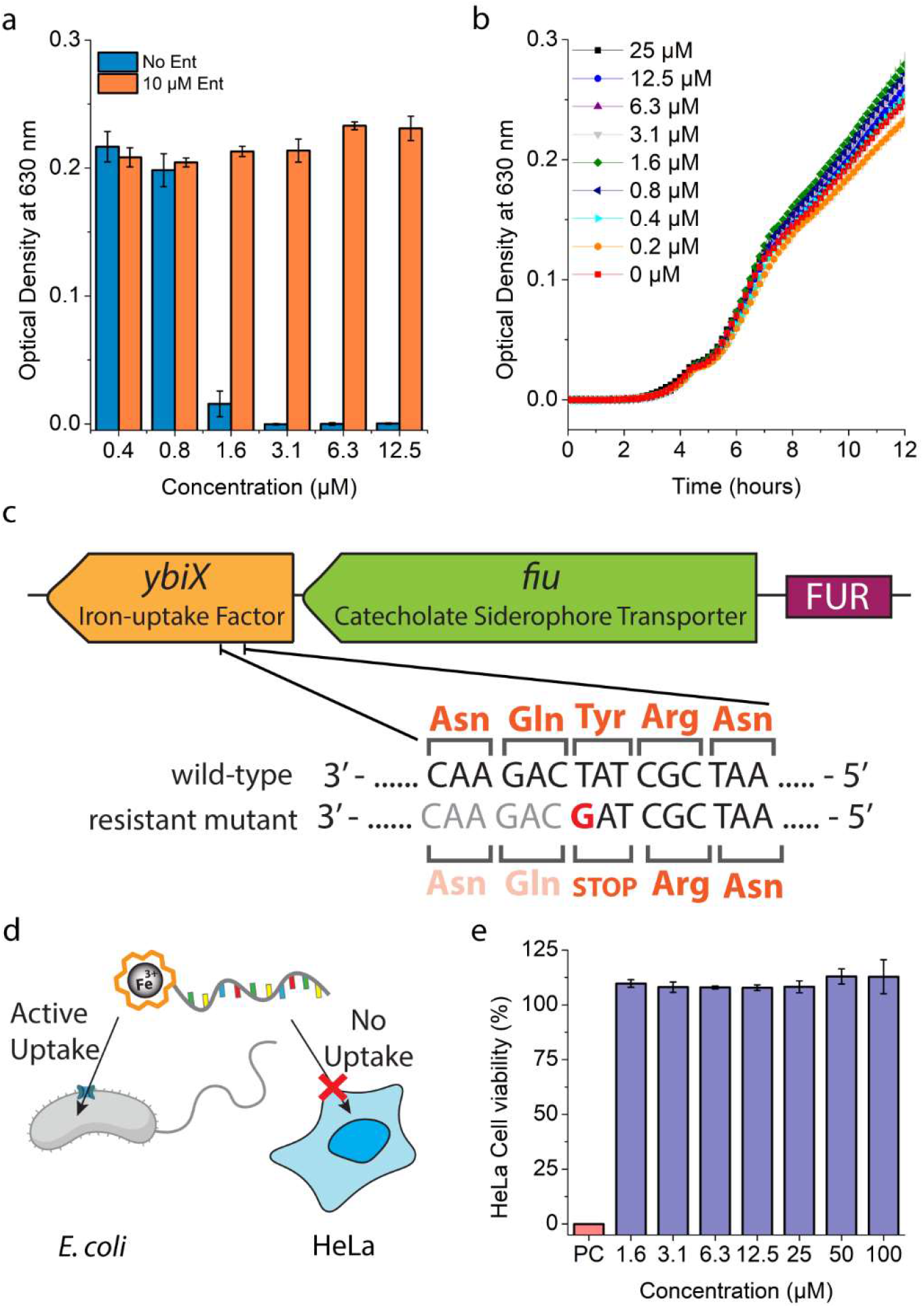
ASO-siderophore conjugate uptake is selective through the iron uptake pathway and non-toxic for human cells. (a) Effect of conjugate (**ASO 2**) uptake due to competition with enterobactin. (b) Growth of the resistant *E. coli* mutant strain in presence of the anti *acpP* PNA-conjugate (**ASO 2**) in iron poor MH2. (c) PacBio whole-genome sequencing of a resistant mutant elucidated an SNP in the *ybiX* gene which results in translation termination. (d) Selective uptake of the ASO-siderophore conjugates. (e) Toxicity of siderophore-PNA conjugate (**ASO 2**) on HeLa cells. PC=10 mM H_2_O_2_. Error-bars represent standard deviations based on three technical replicates.

To further validate that the antibacterial ASOs are transported through the siderophore-mediated iron uptake pathway, we generated resistant *E*.*coli* mutants by prolonged incubation with 6.4 μM of the ASO-siderophore conjugate (**ASO 2**) (4xMIC).^50^ When isolated and regrown, the mutant was not affected by the conjugate even at a concentration 16-fold the MIC of the wildtype (**Fig. 5b**). To elucidate the underlaying mechanism of resistance towards the ASO-siderophore conjugate, the mutant was analyzed by PacBio whole-genome sequencing. One single-nucleotide polymorphism (SNP) was found in the resistant mutant. This T→G mutation in base 838,947 of the *E. coli* K12 genome causes a Tyr→Ter mutation in the *ybiX* gene resulting in premature termination of translation (**Fig. 5c**). The gene encodes a suspected iron-uptake factor^51^ and forms an operon with the *fiu* gene, encoding the outer membrane iron-catecholate transporter.^52^ *ybiX* is a homologue of the *piuC* gene in *Pseudomonas aeruginosa (P. aeruginosa*) that was shown to be involved in siderophore uptake.^34^ Repression of *piuC* in *P. aeruginosa* resulted in resistance towards siderophore-based antibiotics *in vitro*.^53,54^ It is expected that a similar mechanism might cause the observed resistance in the *E. coli* mutant, confirming that ASO-siderophore conjugate uptake is likely mediated through the iron-uptake pathway.

### Toxicity to human cells

Antibacterial ASOs have so far been made mostly with CPPs to enhance bacterial uptake.^12,14,15,55^ CPPs have been shown to transport ASOs into bacteria, but also suffer from toxicity towards human cells, stemming in part from a lack of selectivity towards bacteria.^20,23^ Since the siderophore-transport system is only present in bacteria, our conjugates should not be transported into human cells and not display any toxicity (**Fig. 5d**). To assess the toxicity, HeLA cells were incubated for 24 hours with increasing concentrations of the anti *acpP* PNA-siderophore conjugate (**ASO 2**) at 37°C and 5% CO_2_. After incubation, the cell viability was quantified using a WST-1 assay. As expected, even the highest concentration of 100 μM did not affect cell viability indicating that the conjugates are non-toxic at the tested concentrations, rendering the siderophore-mediated delivery a promising strategy for *in vivo* use.

## Conclusions

In this study, we describe the use of siderophore-mediated delivery of antibacterial ASOs. The conjugation of a synthetic tris-catechol siderophore to a 10-nt PNA ASO, targeting the essential *acpP* gene, resulted in drastically improved antibacterial effect in comparison to the naked PNA (MICs of 1.6 and >100 μM, respectively). The observed antibacterial effect is likely due to binding of the ASO to target mRNA as demonstrated through down regulation of β-galactosidase when targeting the *lacZ* gene. To determine the minimal structural requirements for siderophore-mediated delivery of ASOs to bacterial cells, we constructed an anti *acpP* PNA-conjugate with a single catechol siderophore. Surprisingly, the single catechol group enabled uptake of the ASO, resulting in inhibition of growth. We envision that this feature will find wide applicability in the field of bacteria-targeting ASOs,^8^ given the straightforward synthesis of the mono-catechol siderophore.

To verify that our strategy can be applied to ASOs of varying backbones, we conjugated a 10-nt MPO-based anti *acpP* ASO to the tris-catechol siderophore and determined the antibacterial effect that showed to be potent with a sub micromolar MIC of 0.8 μM. We believe this will be useful because several MPO-based ASOs are FDA approved, indicating safe use in humans.^56^

To confirm the uptake-route of the ASO-siderophore conjugates, we performed a competition assay with the endogenous siderophore enterobactin. The presence of 10 μM enterobactin resulted in a >8 fold increase in MIC, which demonstrates that the ASO-siderophore conjugate is likely taken up via the same route. By exposing *E. coli* to increasing concentrations of the anti *acpP* conjugate, we isolated a resistant mutant strain. PacBio whole genome sequencing revealed an SNP in the *ybiX* gene, which is a suspected iron-uptake factor and forms an operon with *fiu* that encodes an outer membrane iron-catecholate transporter. Together these results show that the ASO-siderophore conjugates are indeed selectively taken up through the iron-transport in bacteria.

Lastly, the *acpP* PNA-siderophore conjugate displayed no toxicity to human HeLa cells up to 100 μM, opening up possibilities for pre-clinical *in vivo* studies.

Taken together, we foresee that the selectivity, simplicity, modularity and low toxicity to human cells makes siderophore-mediated conveyance of ASOs a promising antibacterial strategy, paving the way for next-generation antibiotics.^4^ Furthermore, we expect that the remarkable result that a single catechol group can transfer large (bio)molecules of >3000 Da into bacteria will be useful in biotechnology.

## Acknowledgements

We would like to thank Ing. Frank Nelissen for help with biochemical experimentation and the Biophysical Department at Radboud University and Dr. Hans Heus for generously granting us access to their facilities. This work was supported by an NWO-XS (OCENW.XS21.1.106) grant from the Dutch Research Council to W.A.V. This work has received funding from the European Research Council under the European Union’s Horizon Europe research and innovation programme under grant agreement number 101041938 (RIBOCHEM) to W.A.V.

